# RTICBM-74 is a Brain-Penetrant CB_1_ Receptor Allosteric Modulator that Reduces Alcohol Intake in Rats

**DOI:** 10.1101/2021.09.13.460113

**Authors:** Dennis Lovelock, Thuy Nguyen, Kalynn Van Voorhies, Yanan Zhang, Joyce Besheer

## Abstract

The endocannabinoid system is implicated in the neuronal mechanisms of alcohol use disorder (AUD), with the cannabinoid receptor subtype 1 (CB_1_) representing a promising target for AUD therapeutic interventions. We have previously shown negative allosteric modulators (NAMs) of the CB_1_ receptor attenuated the reinstatement of other drugs of abuse including cocaine and methamphetamine in rats; however, their effects on alcohol-related behaviors have not been investigated. Here, we tested the pharmacokinetic properties of one such CB_1_ NAM, RTICBM-74, and its effects on alcohol self-administration in rats. RTICBM-74 showed low aqueous solubility and high protein binding but had excellent half-life and low clearance against rat liver microsomes and hepatocytes, and excellent brain penetrance in rats. RTICBM-74 pretreatment specifically reduced alcohol intake across a range of doses in male or female Wistar or Long-Evans rats that were trained to self-administer alcohol. These effects were similar to the CB_1_ antagonist/inverse agonist rimonabant which was tested as a positive control. Importantly, RTICBM-74 was effective at reducing alcohol intake at doses that did not affect locomotion or sucrose self-administration. Our findings suggest that CB_1_ NAMs such as RTICBM-74 have promising therapeutic potential in treatment of AUD.

**SIGNIFICANCE STATEMENT:** The present work shows that a metabolically stable and brain penetrant CB_1_ receptor negative allosteric modulator (NAM) reduces alcohol self-administration in rats without affecting locomotion or sucrose self-administration, suggesting potential therapeutic relevance for the treatment of AUD.

## INTRODUCTION

Alcohol use disorder (AUD) is a chronic relapsing brain disease characterized by compulsive alcohol use despite adverse consequences (Koob, 2013). According to the 2015 National Survey on Drug Use and Health (NSDUH), 15.1 million adults ages 18 and older had AUD in the United States. Three medications, naltrexone (opioid antagonist), acamprosate (NMDA receptor agonist), and disulfiram (acetaldehyde dehydrogenase inhibitor), have been approved by the US Food and Drug Administration (FDA) to treat AUD; however, they have only modest efficacy (Heilig and Egli, 2006), illustrating the need for additional therapeutic agents. The cannabinoid type-1 receptor, a key member of the endocannabinoid system, plays a profound role in alcohol related behaviors and has been considered a promising target for alcohol addiction treatment.

The CB_1_ receptor, well known for its role in addiction of multiple drugs (Parsons and Hurd, 2015), contributes to alcohol related behaviors including alcohol consumption, alcohol tolerance, alcohol dependence and addiction. CB_1_ agonist CP55,940 dose-dependently increased ethanol consumption (Gallate et al., 1999), whereas CB_1_ antagonist/inverse agonist rimonabant (SR141716) reduced ethanol intake in a number of preclinical models of ethanol consumption (Arnone et al., 1997; Economidou et al., 2006; Colombo et al., 2007; Hansen et al., 2008; Maccioni et al., 2009). CB_1_ knock-out (KO) mice displayed diminished ethanol-induced conditioned place preference (Houchi et al., 2005). Acute administration of rimonabant completely abolished the alcohol deprivation effect (i.e., relapse-like drinking) and motivation to drink in alcohol-preferring rats (Serra et al., 2002; Colombo et al., 2004; Vinod et al., 2012), while chronic administration during alcohol exposure reduced severity of alcohol withdrawal-induced handling-induced convulsions in both alcohol-preferring and alcohol-avoiding mice (Vinod et al., 2008). Finally, rimonabant treatment prior to a reinstatement session reduced cue-induced relapse (Cippitelli et al., 2005; Economidou et al., 2006; Economidou et al., 2007). Together, these results demonstrate the promise of CB_1_ blockade as potential treatments of AUD. Unfortunately, after its initial approval in Europe for the treatment of obesity (Cahill and Ussher, 2007), rimonabant was eventually withdrawn from the market because of untoward side effects associated with its use in humans including depression, anxiety or even suicidal ideation (Christensen et al., 2007; Moreira et al., 2009; Beyer et al., 2010).

Several alternate approaches to modulate the CB1 signaling that may avoid the psychiatric side-effects of CB1 antagonists/inverse agonists have been pursued (Nguyen et al., 2019; Cinar et al., 2020), among which allosteric modulation has received particular attention. Allosteric modulators of G protein-coupled receptors (GPCRs) are molecules that bind at topographically distinct allosteric sites and either potentiate (positive allosteric modulators or PAMs) or inhibit (negative allosteric modulators or NAMs) the binding and/or signaling of an orthosteric ligand (Kenakin and Miller, 2010). Several CB_1_ NAMs have been reported thus far (Nguyen et al., 2017b), including the two most extensively studied Org27569 and PSNCBAM-1. These NAMs were able to attenuate or block functional responses of CB_1_ agonists such as CP55940 in several in vitro assays (Price et al., 2005; Horswill et al., 2007). We found that Org27569 and PSNCBAM-1, like rimonabant, resulted in a dose-related attenuation of cue- and/or drug-induced reinstatement of cocaine- and/or methamphetamine-seeking behavior (Jing et al., 2014; Nguyen et al., 2017a). Continued efforts from our team resulted in the identification of CB_1_ NAM RTICBM-74, which displayed improved metabolic stability over PSNCBAM-1. Despite their similar in vitro potency, RTICBM-74 at 10 mg/kg produced the same degree of attenuation as that of 30 mg/kg of PSNCBAM-1 in prime-induced reinstatement of cocaine seeking behavior, suggesting improved in vivo potency (Nguyen et al., 2017a). However, the effects of RTICBM-74 on alcohol reinforcement have yet to be studied.

Here, we first assessed the ADME (absorption, distribution, metabolism, and excretion) and pharmacokinetic properties of RTICBM-74 and then investigated its effects on alcohol self-administration in rats. To assess the reinforcer specificity of RTICBM-74, we also tested sucrose self-administration in parallel and tested rimonabant as a positive control.

## MATERIALS AND METHODS

### Drugs

RTICBM-74 was synthesized at Research Triangle Institute following previously published procedures (purity >98%) (Nguyen et al., 2017a). For behavioral experiments RTICBM-74 was suspended in a mixture of saline, tween-80 and DMSO of an 8:1:1 ratio. Rimonabant (National Institute on Drug Abuse Drug Supply Program) was dissolved in saline and 2 drops of Tween-80 per 5 ml of saline. All injections were administered at a volume of 1 mL/kg.

### RTICBM-74 properties

#### Receptor Selectivity

The off-target selectivity of RTICBM-74 was assessed in radioligand binding assays against a panel of ∼50 GPCRs, ion channels, and transporters at a single concentration of 10 μM by NIMH Psychoactive Drug Screening Program (PDSP) according to their standard protocols. These targets included adrenergic (α_1A_, α_1B_, α_1D_, α_2A_, α_2B_, α_2C_, β1, β2, β3), dopamine (D1 - D5), GABA_A_, histamine (H1 - H4), muscarinic (M1 - M5), opioid (μ, κ, δ), serotonin (5-HT1A, 5-HT1B, 5-HT1D, 5-HT1E, 5-HT2A, 5-HT2B, 5-HT2C, 5-HT3, 5-HT5A, 5-HT6, 5-HT7A), sigma (σ1, σ2), dopamine transporter (DAT), norepinephrine transporter (NET), serotonin transporter (SERT), and BZP Rat Brain Site. The percentage of inhibition was the average of two determinations. When significant displacement of radioligand was observed (>50% inhibition at 10 μM), complete concentration-dependent displacement curves (in duplicate) were constructed to generate IC_50_ values. IC_50_ values were determined by a nonlinear regression analysis. The equilibrium dissociation constant (Ki) was calculated with the Cheng–Prusoff equation using the observed IC_50_ of RTICBM-74, the concentration of radioligand, and the historical values of *K*_d_ of the ligand. The full methods and references can be found at: http://pdspdb.unc.edu/pdspWeb.

### ADME Studies

#### Hepatocyte Stability

Plateable cryopreserved rat hepatocytes (BioreclamationIVT, USA) were thawed in recovery media and diluted in plating media to a viable cell density of 1×10^6^ cells/mL. Viability was determined by trypan blue exclusion. A 50 μL suspension of cells was added to each well of a 96-well plate and pre-incubated for 10 minutes at 37°C, 5% CO_2_. At time zero, 50 μL of test articles and reference compounds (testosterone and ethoxycoumarin) were added in duplicate to reach a final concentration of 1 μM and to initiate the reaction. At selected time points (15, 30, 60, 120 and 240 min), the plate was removed from the incubator and the wells were quenched with 150 μL of (75/25) ice-cold acetonitrile (ACN) in water containing 2 % formic acid and internal standard (0.1μM glyburide and labetalol). The 0 min time points were spiked in the pre-quenched wells. Samples were then centrifuged and supernatants were transferred to a new plate for analysis by LC-MS/MS and remaining parent drug was used to determine the half-life and intrinsic clearance using Microsoft Excel.

#### Kinetic Solubility

The kinetic solubility of the test compound was measured in commercial PBS, which consisted of potassium phosphate monobasic 1 mM, sodium phosphate dibasic 3 mM and sodium chloride 155 mM. A 10 μL of test compound stock solution (20 mM DMSO) was combined with 490 μL of PBS buffer to reach a targeted concentration of 400 μM. The solution was agitated on a VX-2500 multi-tube vortexer (VWR) for 2 hours at room temperature. Following agitation, the sample was filtrated on a glass-fiber filter (1 μm) and the eluate was diluted 400-fold with a mixture of acetonitrile: water (1:1). On each experimental occasion, nicardipine and imipramine were assessed as reference compounds for low and high solubility, respectively. All samples were assessed in triplicate and analyzed by LC-MS/MS using electrospray ionization against standards prepared in the same matrix.

#### Plasma Protein Binding

Test compound and controls (1 μM) were spiked in rat plasma (BioreclamationIVT, Westbury, NY, USA) and aliquoted in triplicate in a high throughput dialysis (HTD) 96-well plate, where the plasma and dialysate buffer were separated by a semi-permeable cellulose membrane (12-14K MWCO). Once sealed, the HTD plate was incubated at 37°C and kept under light agitation for 6 hours, until equilibrium was reached. Plasma and buffer samples were then extracted along with their corresponding standard curve samples using ice-cold acetonitrile in methanol (1:1). After centrifugation, supernatants from both plasma- and buffer-containing samples were further diluted prior to be submitted to bioanalysis by LC-MS/MS. Acebutolol and warfarin served as low-bound and highly-bound controls, respectively. Determination of the percentage of plasma protein binding and recovery was achieved in Microsoft Excel.

#### Bidirectional MDCK-mdr1 permeability

MCK-mdr1 cells were seeded onto permeable polycarbonate supports in 12-well Costar Transwell plates and allowed to grow and differentiate for 3 days. On day 3, culture medium (DMEM supplemented with 10% FBS) was removed from both sides of the transwell inserts and cells were rinsed with warm HBSS. After the rinse step, the chambers were filled with warm transport buffer (HBSS containing 10 mM HEPES, 0.25% BSA, pH 7.4) and the plates were incubated at 37oC for 30 min prior to TEER (Trans Epithelial Electric Resistance) measurements. The buffer in the donor chamber (apical side for A-to-B assay, basolateral side for B-to-A assay) was removed and replaced with the working solution (10 μM test article in transport buffer). The plates were then placed at 37°C under light agitation. At designated time points (30, 60 and 90 min), an aliquot of transport buffer from the receiver chamber was removed and replenished with fresh transport buffer. Samples were quenched with ice-cold ACN containing internal standard and then centrifuged to pellet protein. Resulting supernatants are further diluted with 50/50 ACN/H2O (H2O only for Atenolol) and submitted for LC-MS/MS analysis. Reported apparent permeability (P_app_) values were calculated from single determination. Atenolol and propranolol were tested as low and moderate permeability references. Bidirectional transport of digoxin was assessed to demonstrate P-gp activity/expression.

#### PK Studies

A PK study of RTICBM-74 was performed using male Sprague-Dawley rats (Paraza Pharma Inc., Montreal, Canada). RTICBM-74 was suspended in 5% Cremorphor, 5% ethanol and 90% saline. On the morning of the PK study, animals were weighed and dosing formulation volumes were calculated accordingly. The compound was injected intraperitoneally (i.p.) to all animals at 10 mg/kg. At selected time points (0.5, 1, 4, 8, and 24 h post-dose), two animals were anesthetized to perform a cardiac puncture to collect blood for plasma analysis, followed by whole body perfusion with phosphate saline buffer (pH 7.4) to wash out any remaining blood from the organs. Brains were harvested and homogenized by polytron 1:4 (w/v) in 25% 2-propanol in water. Brain homogenates were extracted for drug quantification of LC-MS/MS.

### Self-administration Experiments

#### Subjects

Adult male Wistar rats (Jackson Laboratories) and adult male and female Long Evans rats (Envigo) were used in this study. Rats were double housed under a 12h light/dark cycle (7am/pm). All experiments were conducted during the light cycle. Animals were continuously monitored and cared for by the veterinary staff of the UNC-Chapel Hill Division of Comparative Medicine. All procedures were carried out in accordance with the NIH Guide for Care and Use of Laboratory Animals and institutional guidelines. All protocols were approved by the UNC Institutional Animal Care and Use Committee (IACUC). UNC-Chapel Hill is accredited by the Association for Assessment and Accreditation of Laboratory Animal Care (AAALAC).

#### Apparatus

Self-administration was conducted in operant chambers (Med Associates, Georgia, VT) located within sound-attenuating cabinets equipped with an exhaust fan to provide ventilation and mask outside noise. Chambers were equipped with two retractable levers on opposite sides of the chamber (left and right), and a cue light was located above each lever. When the response requirement was met on the left (active) lever, a cue light (directly above the lever) and a stimulus tone were presented for the duration of the alcohol reinforcer delivery (0.1mL of solution into a well on the left side of the chamber across 1.66s). Responding during reinforcer delivery and on the right (inactive) lever was recorded, but had no programmed consequences. Chambers were also equipped with 4 parallel infrared beams across the bar floor to measure general locomotor activity throughout the session. The number of beam breaks for the entire session was collected and this total was divided by the session length (30 min) to represent the locomotor rate (beam breaks/min).

### Alcohol self-administration training

Rats were trained to self-administer a 15% (v/v) alcohol on a fixed ratio 2 (FR2) schedule of reinforcement in 30-minute sessions, five days a week (M-F) via sucrose fading as described in (Randall et al., 2017; Randall et al., 2019; Makhijani et al., 2020; Ornelas et al., 2021). Sucrose fading began with self-administration of 10% sucrose and 2% alcohol (2A/10S), then 5A/10S, 10A/10S, 10A/5S, 15A/5S, 15A/2S on subsequent sessions, and then remained at 15A for the duration of training. Rats had approximately 4 months of self-administration training prior to testing and rats in SA2 and SA3 were used in an unrelated drug test study that concluded a week prior to the initiation of this study.

### Self-administration Experiment 1 (SA1): Effect of RTICBM-74 on alcohol and sucrose self-administration in female Long-Evans rats

To measure the effect of RTICBM-74 on maintenance of alcohol and sucrose self-administration, female Long-Evans rats received RTICBM-74 (0, 5, 7.5, 10 mg/kg, IP) 20 minutes prior to a self-administration session. A within subject design was used such that each rat (n = 11) received each treatment in a random order. Test days were on Tuesdays and Thursdays with standard self-administration sessions on Monday, Wednesday and Friday. Alcohol lever responses had to be at least 80% of baseline self-administration (average responding in the 2 sessions preceding the test) for a rat to be tested and all rats met this criterion. 1 week after conclusion of testing, the same cohort of rats was trained to self-administer 0.8% (w/v) sucrose solution on a FR2 schedule. Rats (n = 11) received the same doses of RTICBM-74 using the same testing procedure and within subject design as when tested with alcohol self-administration.

### SA2: Effect of RTICBM-74 on alcohol self-administration in male Wistar rats

To measure the effect of RTICBM-74 on maintenance of alcohol self-administration in different strains or sexes, male Wistar rats received RTICBM-74 (0, 7.5, 10 mg/kg, IP) 20 minutes prior to a self-administration session. A within subject design was used such that each rat (n = 11) received each treatment in a random order. The testing schedule was the same as described in Experiment 1 (SA1).

### SA3: Effect of RTICBM-74 on sucrose self-administration in male Wistar rats

A separate cohort of male Wistar rats (n=11) were trained to self-administer 0.5% (w/v) sucrose solution on an FR2 schedule and received the same doses and treatment schedule of RTICBM-74 as described in Experiment 2. A lower concentration of sucrose was used compared to Experiment 1 in order to more closely match total lever presses between alcohol (SA2) and this sucrose self-administration group.

### SA4: Effect of rimonabant on alcohol self-administration in Long-Evans rats

To assess the effects of rimonabant on alcohol self-administration, male and female Long-Evans rats received rimonabant (0, 1, 3 mg/kg, IP) 30 minutes prior to an alcohol self-administration session. A within subject design was used such that each rat (n = 12/sex) received each treatment in a random order. The testing schedule was the same as described in Experiment 1.

#### Self-administration data analysis

Alcohol or sucrose lever responses, inactive lever responses, alcohol or sucrose intake (g/kg or mL/kg, respectively; estimated from the number of reinforcers received) and locomotor rate were analyzed with one-way repeated measures analysis of variance (RM-ANOVA) with RTICBM-74 dose as a factor. Alcohol and sucrose intake data for all experiments, as calculated from number of reinforcers received and body weight, are included in Table 3. Cumulative alcohol lever responses across the session were analyzed by two-way RM-ANOVA with RTICBM-74 dose and session time as factors. In all cases, post-hoc analysis (Tukey) was used to determine specific differences from the vehicle condition. For all analyses, significance was set at p ≤ 0.05.

**Table 1.** ADME properties of RTICBM-74

| Rat Liver Microsome<sup>a</sup> | | Hepatocyte | | Kinetic solubility ($\mu\text{M}$) | %Protein bound | MDCK-mdr1 | | |
| --- | --- | --- | --- | --- | --- | --- | --- | --- |
| $t_{1/2}$ (min) | $Cl_{\text{int}}$ (mL/min/kg) | $t_{1/2}$ (min) | $Cl_{\text{int}}$ (mL/min/kg) | | | $P_{\text{app}}$ (A to B) ($10^{-6}$ cm/s) | $P_{\text{app}}$ (B to A) ($10^{-6}$ cm/s) | efflux ratio |
| >300 | <4.6 | 461 | 14.5 | 1.09 | 99.9% | 0.3 | 0.2 | 0.7 |

<sup>a</sup> Reported previously in (Nguyen et al., 2017a)

**Table 2.** Pharmacokinetic analysis of RTICBM-74 in rats (i.p. 10 mg/kg)

| | $C_{\text{max}}$ (ng/mL) | $t_{\text{max}}$ (h) | $\text{CL}_{\text{F}}$ (mL/min/kg) | $\text{AUC}_{\text{inf}}$ (ng/mL*h) |
| --- | --- | --- | --- | --- |
| **Plasma** | 70.2 | 6.0 | 119.4 | 1476 |
| **Brain** | 272.0 | 4.5 | 68.4 | 2481 |
| **Brain/plasma ratio** |  |  |  | 1.7 |

**Table 3.**
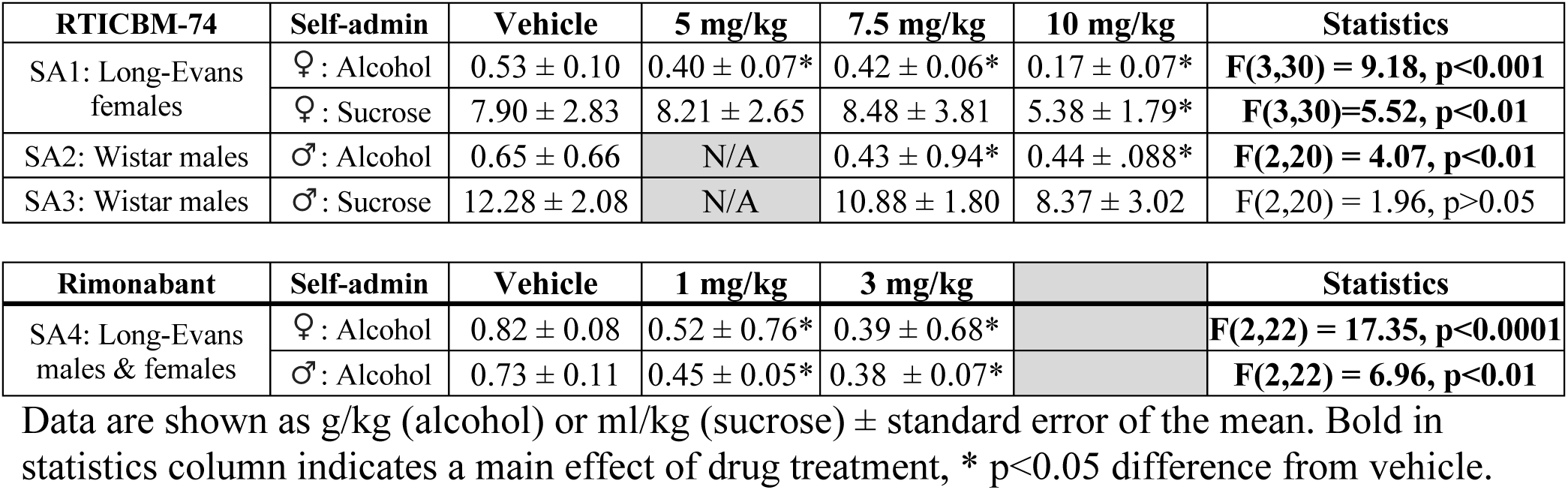
Alcohol intake from self-administration experiments. Data are shown as g/kg (alcohol) or ml/kg (sucrose) ± standard error of the mean. Bold in statistics column indicates a main effect of drug treatment, * p<0.05 difference from vehicle.

| **RTICBM-74** | **Self-admin** | **Vehicle** | **5 mg/kg** | **7.5 mg/kg** | **10 mg/kg** | **Statistics** |
| --- | --- | --- | --- | --- | --- | --- |
| SA1: Long-Evans females | ♀ : Alcohol | 0.53 ± 0.10 | 0.40 ± 0.07* | 0.42 ± 0.06* | 0.17 ± 0.07* | **F(3,30) = 9.18, p<0.001** |
|  | ♀ : Sucrose | 7.90 ± 2.83 | 8.21 ± 2.65 | 8.48 ± 3.81 | 5.38 ± 1.79* | **F(3,30)=5.52, p<0.01** |
| SA2: Wistar males | ♂ : Alcohol | 0.65 ± 0.66 | N/A | 0.43 ± 0.94* | 0.44 ± .088* | **F(2,20) = 4.07, p<0.01** |
| SA3: Wistar males | ♂ : Sucrose | 12.28 ± 2.08 | N/A | 10.88 ± 1.80 | 8.37 ± 3.02 | F(2,20) = 1.96, p>0.05 |

| **Rimonabant** | **Self-admin** | **Vehicle** | **1 mg/kg** | **3 mg/kg** |  | **Statistics** |
| --- | --- | --- | --- | --- | --- | --- |
| SA4: Long-Evans males & females | ♀ : Alcohol | 0.82 ± 0.08 | 0.52 ± 0.76* | 0.39 ± 0.68* |  | **F(2,22) = 17.35, p<0.0001** |
|  | ♂ : Alcohol | 0.73 ± 0.11 | 0.45 ± 0.05* | 0.38 ± 0.07* |  | **F(2,22) = 6.96, p<0.01** |

## RESULTS

### ADME Studies

The metabolic stability of RTICBM-74 in rat liver microsomes has previously been reported by us (Nguyen et al., 2017a). In hepatocytes, RTICBM-74 showed a clearance of 14.5 mL/min/kg, with a half-life (t_1/2_) of 461 minutes (Table 1). RTICBM-74 had kinetic solubility of 1.09 μM in the phosphate-buffered saline (PBS) solution at pH = 7.4. When tested using the MDCK-mdr1 cell line expressing the human P-glycoprotein efflux transporter, RTICBM-74 showed apparent permeability P_app_ values of 0.3 (A-to-B) and 0.2 10^−6^ cm/s (B-to-A), respectively. The efflux ratio was 0.7.

#### Receptor Selectivity

RTICBM-74 at 10 μM showed <50% inhibition of all the GPCRs, enzymes or transporters tested.

#### PK Study

Following an intraperitoneal (i.p.) dose of 10 mg/kg in rats, RTICBM-74 reached peak plasma concentration (C_max_ = 70 ng/mL) at 6 h post-dose. The brain concentration peaked at 4.5 h with a brain C_max_ of 272 ng/mL. Higher concentrations in the brain than plasma were observed at all data points tested (0.5, 1, 4, 8, 24h). RTICBM-74 was eliminated from the brain with a clearance of 68.4 mL/min/kg and from the plasma with a clearance of 119.4 mL/min/kg. The overall brain to plasma ratio (K_p_) was 1.7, as determined by the AUC_inf_ ratio of brain and plasma.

### Self-administration experiments

#### SA1: RTICBM-74 and alcohol self-administration in female Long-Evans rats

A one-way RM ANOVA found a main effect of RTICBM-74 on alcohol lever responses [F(3, 30)=7.90, p<0.0001] with significant reductions at all three doses (p’s <0.05; figure 1A). Across 5 minutes bins (figure 1B), there were main effects of dose [F(5,50)=81.73, p<0.0001] and time bin [F(3,30)=7.37, p<0.001] as well as an interaction [F(15,150)=5.43, p<0.0001]. Post-hoc comparisons revealed that the 7.5 and 10 mg/kg doses reduced responding in the first 5 minutes (p<0.01, p<0.0001), while all three doses reduced active lever responses from the second time bin onward (p’s<0.0001). In locomotor activity there was an effect of treatment [F(3,30)=14.60, p<0.0001] with a reduction found only with the highest dose (p<0.0001; figure 1C), indicating a possible motor impairment at the highest dose. This finding suggests that the reduction in self-administration at this 10 mg/kg dose may be a nonspecific reduction related to a motor impairment.

**Figure 1:**
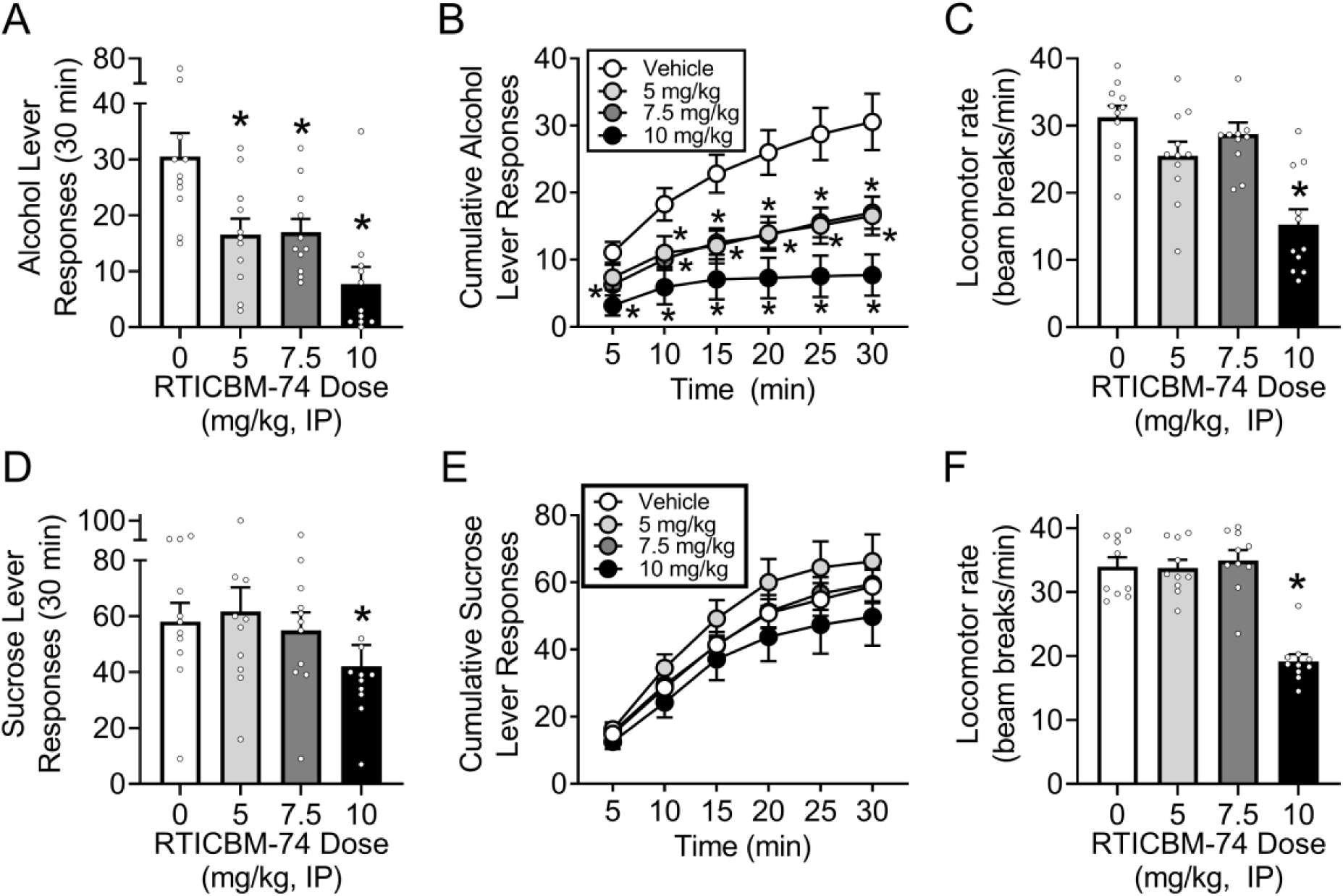
Experiment SA1 found that RTICBM-74 reduced alcohol lever responses in female Long-Evans rats at all doses tested (A,B) while locomotor rate was only reduced by the highest dose (C). Only the highest dose reduced lever responding for a sucrose reward (D,E), which is likely explained by the generalized reduction in motor activity at this dose (F). *p<0.05

#### SA1: RTICBM-74 and sucrose self-administration in female Long-Evans rats

In sucrose self-administration there was a main effect of drug treatment [F(3,30)=5.19, p<0.01] with a reduction found only in the 10 mg/kg group (p<0.05; figure 1D). A similar result was found with locomotor activity [F(3,30)=65.46, p<0.01] as again a reduction was found with treatment of the highest dose (p<0.0001; figure 1F), confirming a likely motor impairment at this dose. Further, the lack of a reduction of sucrose self-administration at the 5 or 7.5 mg/kg dose suggests that the reductions in alcohol self-administration at those doses are specific to alcohol reinforcement.

#### SA2: RTICBM-74 and alcohol self-administration in male Wistar rats

A one-way RM ANOVA found that RTICBM-74 reduced lever responses [F(2, 20)=4.79, p<0.05] at the 7.5 and 10 mg/kg doses (p’s < 0.05; figure 2A). Across time (figure 2B) there was a main effect of time bin [F(5,165)=64.83, p<0.0001], a trend for an effect of dose (p=0.085), and a significant interaction [F(10,65)=2.41, p<0.05]. Post-hoc comparisons revealed that the 7.5 mg/kg dose was lower than vehicle at the 15 min time bin (p<0.05), while the 10 mg/kg dose reduced responding at 20, 25, and 30 min (p’s<0.05). RTICBM-74 did not affect locomotor rate but approached a statistical trend (p=0.079; figure 2C).

**Figure 2:**
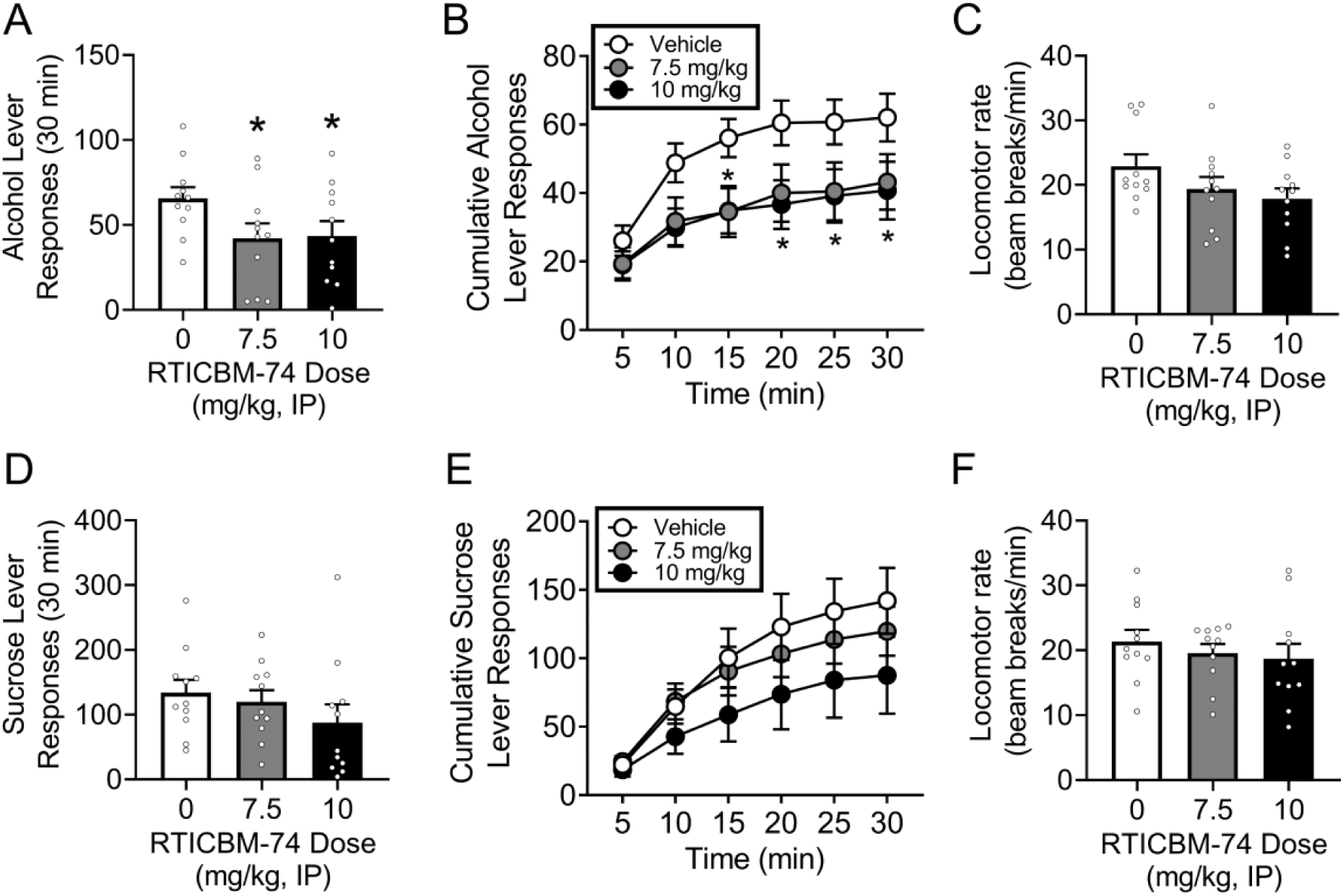
In Experiment SA2, RTICBM-74 reduced lever responses for alcohol in Male Wistar rats (A,B) and had no effect on locomotor rate during the test session (C). Experiment SA3 examined the effect of RTICBM-74 on self-administration for sucrose finding no effects (D-F), suggesting the results from Experiment SA2 are reinforcer-specific. *p<0.05

#### SA3: RTICBM-74 and sucrose self-administration in male Wistar rats

RTICBM-74 had no significant effects on behaviors in sucrose self-administration or locomotion (figures 2D-F), suggesting specificity of the reduction to the alcohol reinforcer.

#### SA4: Rimonabant self-administration in male and female Long Evans rats

In males, rimonabant reduced alcohol lever responding [F(2, 22)=6.22, p<0.01] at 1 and 3 mg/kg (p’s<0.05; figure 3A). A two-way RM ANOVA analyzing lever responses in 5 minute bins (figure 3B) found main effects of rimonabant [F(2,33)=5.35, p<0.01] and time bin [F(5,165)=44.23, p<0.0001] with a significant interaction [F(10,165)=2.33, p<0.05]. Post-hoc analyses indicated that both doses of rimonabant reduced the total number of lever responses starting with the third time bin (p’s < 0.05). In females, rimonabant also reduced lever pressing for alcohol [F(2,22)=15.13, p<0.0001] at the 1 (p<0.01) and 3 (p<0.0001) mg/kg doses (figure 3D). When analyzed across the session (figure 3E) there were main effects of rimonabant [F(2,22)=18.06, p<0.0001], time bin [F(5,55)=21.66, p<0.0001], and a significant interaction [F(10,110)=2.63, p<0.01]. Post-hoc comparisons found that both doses of rimonabant reduced lever pressing at every time bin compared to vehicle (p’s<0.01]. There was a main effect of rimonabant on locomotor activity [F(2,22)=8.92, p<0.01] where the 3 mg/kg dose increased the number of beam breaks/min (p<0.01; figure 3F). In sum, rimonabant pretreatment, which was used as a positive control, induced reductions in alcohol self-administration in males and females consistent with results previously found in males (Freedland et al., 2001; Colombo et al., 2007; Maccioni et al., 2009).

**Figure 3:**
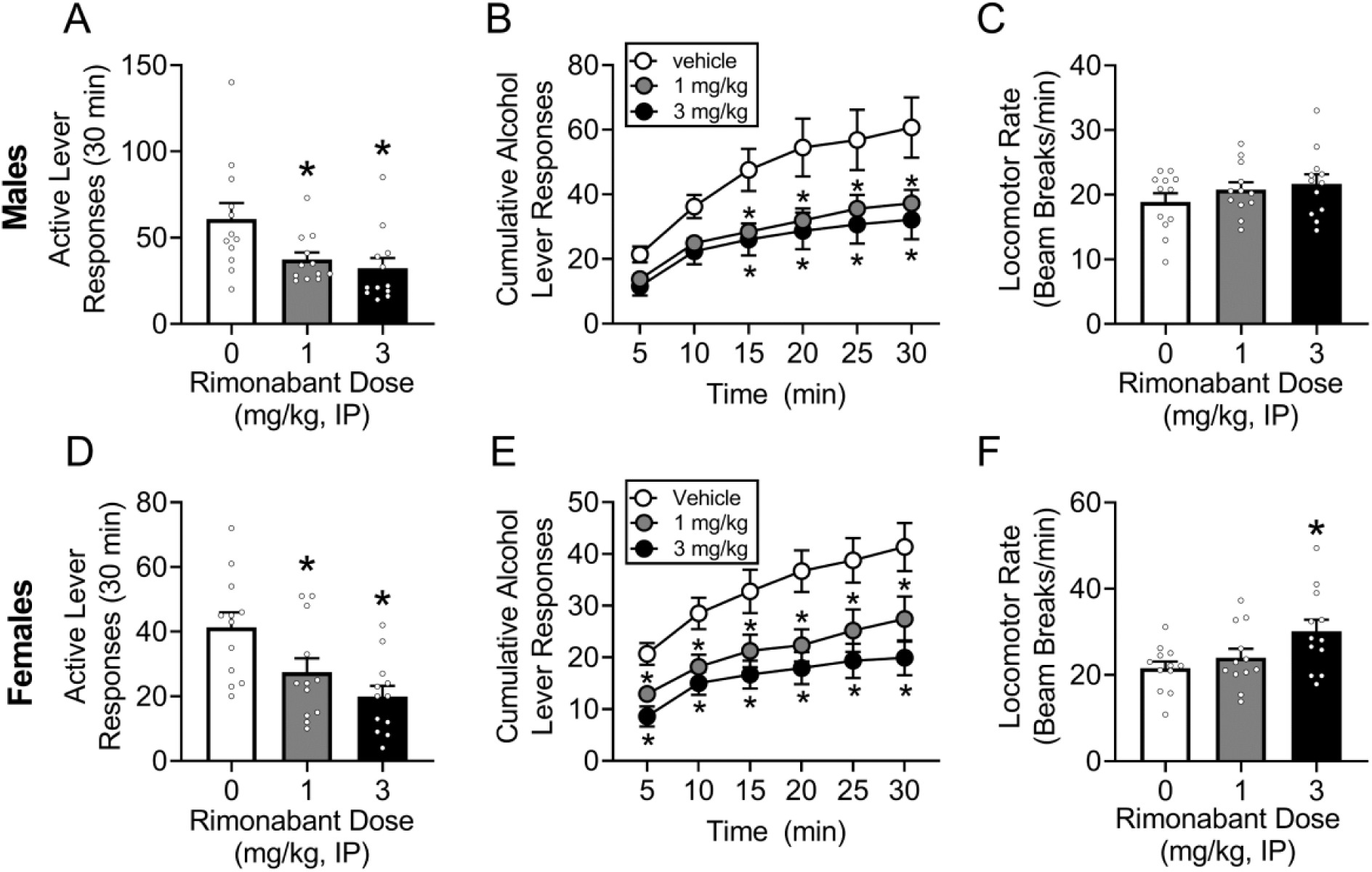
Experiment SA4 tested rimonabant as a positive control experiment in male and female Long-Evans rats. As expected, rimonabant reduced lever responding in both males (A,B) and females (D,E). The high dose of rimonabant increased locomotor activity during the test session in females (F), though this did not result in a concordant increase in lever responses.*p<0.05

## DISCUSSION

The present report evaluated the pharmacokinetic and behavioral characteristics of the potent CB_1_ NAM, RTICBM-74. RTICBM-74 showed good in vitro potency at the CB_1_ receptor, with IC_50_ values of 23 nM in calcium mobilization assay and 153 nM in [^35^S]GTPγS assay (Nguyen et al., 2017a). In rat liver microsomes, RTICBM-74 displayed excellent metabolic stability with long half-life (>300 min) and low clearance (<4.6 μL/min/mg) (Nguyen et al., 2017a). In hepatocytes, similar high stability was observed with RTICBM-74, with half-life of 461 min and low clearance (14.5 μL/min/mg). However, poor aqueous solubility (1.09 μM) and high plasma protein binding (99.9%) were observed. These are consistent with the high lipophilicity and low solubility of many cannabinoid ligands including rimonabant (solubility of 2.6 μM at pH 7.4) (Alelyunas et al., 2010). The high protein binding may have, in part, contributed to the observed good metabolic stability of RTICBM-74, with only small fractions of the compounds being unbound and to be metabolized. This low unbound drug concentration may have also resulted in the modest apparent permeability values (P_app_ of 0.3 (A-to-B) and 0.2 × 10^−6^ cm/s (B-to-A)) observed in the MDCK-mdr1 assay. The efflux ratio of 0.7 suggests that RTICBM-74 was not a P-glycoprotein substrate.

The low P_app_ values in the MDCK-mdr1 assay may suggest low brain penetration for RTICBM-74; however, higher concentrations in brain than plasma at all time points and a significant higher brain C_max_ than plasma (272.0 ng/mL vs. 70.2 ng/mL) were observed in the PK study in rats. In addition, a brain/plasma ratio (K_p_ = AUC_brain_/AUC_plasma_) of 1.7 was obtained. These results support good brain penetration for RTICBM-74. However, the high T_max_ of 4.5h in the brain in the PK studies and low apparent permeability (P_app_) in the MDCK-mdr1 assay together suggest that RTICBM-74 is slow in entering the brain. This is again likely because of the high protein binding of RTICBM-74 and the resulting low unbound drug levels in both plasma and brain. RTICBM-74 was cleared from the brain and plasma at moderate or high rate at 68.4 and 119.4 mL/min/kg, respectively. Together, these results indicate RTICBM-74 is a brain-penetrant CB_1_ allosteric modulator, albeit with low penetration rate.

RTICBM-74 was administered 30 minutes prior to alcohol self-administration attenuated alcohol intake. All three doses tested (5, 7.5, 10 mg/kg) reduced alcohol self-administration in Long-Evans female rats (figure 1A), and both the highest doses reduced initial intake in the first 5 minute of the session (figure 1B), which could be interpreted as a reduction in initiation to drink. However, once the session progressed and alcohol was consumed, a greater reduction in ongoing alcohol self-administration was observed, suggesting that RTICBM-74 is more effective at reducing ongoing drinking. General locomotor activity was decreased only at the highest dose which may have contributed to the reduction in self-administration at that dose (figure 1C). However, as locomotor activity was not reduced at lower doses (5 and 7.5 mg/kg) the reduced consumption was likely not due to a non-specific motor reduction. The effects of RTICBM-74 were tested on sucrose self-administration to test for reinforcer specificity, and a reduction in sucrose intake was only seen at the highest dose (10 mg/kg; figure 1D), which again induced a concordant reduction in locomotor activity that suggests general motor impairments (figure 1F).

To examine replicability and generalizability of these effects across sex and strain, we tested the two highest doses of RTICBM-74 using the same self-administration procedures in male Wistar rats. Again, all doses reduced alcohol self-administration (figure 2A). In male Wistar rats the effect was not immediately detected but emerged after 15 (10 mg/kg) or 20 (7.5 mg/kg) minutes of self-administration (figure 2B), again supporting that RTICBM-74 is more effective at reducing ongoing drinking. Notably, there was no reduction in locomotor activity at the highest dose (10 mg/kg) as was seen in the Long-Evans females (figure 2C), suggesting species differences. Likewise, RTICBM-74 had no effect on sucrose self-administration (figure 2D-F). The Wistar male rats were less sensitive to the locomotor reduction at the highest dose as compared to the Long-Evans females, but notably the 7.5 mg/kg dose effectively and specifically reduced alcohol self-administration in both groups.

Rimonabant has been shown to reduce operant alcohol-self administration across rat strains (Freedland et al., 2001; Colombo et al., 2004; Hansson et al., 2007; Malinen and Hyytiä, 2008). We therefore tested rimonabant as the positive control experiment on alcohol self-administration. As expected, both low and high (1 and 3 mg/kg) doses of rimonabant effectively reduced alcohol lever responses (figure 3A,D) in male and female Long Evans rats. Across time a similar pattern was seen as in prior experiments where the reduction became significant over time in males but was immediately apparent in females (figure 3 B,E). Rimonabant did not reduce locomotor activity at either dose, although the high dose increased locomotor activity in females. This increase was unexpected as few studies have tested females, but the increase in generalized activity did not affect the expected reduction in alcohol intake. Overall, we found that RTICBM-74 effectively reduced alcohol intake as effectively as rimonabant across multiple rat strains and in both sexes, and showed that the reduction effect was specific to alcohol.

In summary, the present studies were designed to characterize RTICBM-74 towards the goal of developing a novel CB_1_ NAM-based treatment for AUD. RTICBM-74 was found to have good *in vitro* potency at the CB_1_ receptor, excellent metabolic stability, and effective CNS penetration, all of which suggest efficacy as a centrally-acting peripherally-administered treatment. IP administration of RTICBM-74 effectively and specifically reduced alcohol self-administration in both male and female rats across multiple strains, without affecting locomotor activity or sucrose intake. Altogether, the findings suggest that CB_1_ NAMs such as RTICBM-74 may have therapeutic potential in treatment of AUD.

## Abbreviations

ACN: acetonitrile
ADME: absorption, distribution, metabolism and excretion
AUC: area under the curve
AUD: alcohol use disorder
CB_1_: cannabinoid receptor subtype 1
GPCR: G-coupled protein receptor
HTD: high throughput dialysis
MDCK: Madin-Darby canine kidney
NAM: negative allosteric modulator
PAM: positive allosteric modulator
Papp: apparent permeability
PK: pharmacokinetics
TEER: Trans Epithelial Electric Resistance

## Acknowledgements

This work was supported in part by the National Institute of Health AA011605 (JB), AA026537 (JB) and DA040693 (YZ) and by the Bowles Center for Alcohol Studies. We are grateful to the NIMH Psychoactive Drug Screening Program (PDSP) for the receptor selectivity testing and to Brayden Fortino for technical assistance with the self-administration studies.

